# Axial Patterning Beyond the Individual: Colony-level Organization in a Siphonophore Colony

**DOI:** 10.64898/2026.04.01.715818

**Authors:** Kohei Oguchi, Akifumi Yao, Hisanori Kohtsuka, Shigeru Kuratani, Toru Miura

**Affiliations:** Misaki Marine Biological Station, School of Science, The University of Tokyo, 1024 Koajiro, Misaki, Miura, Kanagawa 238-0225, Japan; Molecular Life History Laboratory, National Institute of Genetics, 1111 Yata, Mishima, Shizuoka 411-8540, Japan; RIKEN Center for Biosystems Dynamics Research, 2-2-3 Minatojima-minami, Chuo, Kobe, Hyogo 650-0047, Japan; Institute of Science Tokyo, Faculty of Dentistry, 2 Chome-12-1 Ookayama, Meguro City, Tokyo 152-8550, Japan

**Author notes:** Corresponding author: Kohei Oguchi.

**Keywords:** coloniality, superorganism, axial patterning, ora-aboral axis, Hox genes, Wnt signaling, zooid differentiation, growth zones

## Abstract

Colonial animals composed of clonally produced units can achieve a high degree of functional integration, challenging the distinction between an individual and a higher-order organism. Siphonophores (Cnidaria: Hydrozoa) exemplify this condition, forming highly organized colonies in which genetically identical zooids are specialized for functions such as locomotion, feeding, and reproduction, and are precisely arranged along a shared stem. All zooids arise from two spatially separated budding zones, the nectosomal and siphosomal growth zones, suggesting that positional information along the stem patterns colony organization at the level of the colony rather than individual zooids. However, the molecular basis of this colony-level axial patterning remains poorly understood. Here, we analyze gene expression along the stem of the siphonophore *Agalma okenii* using RNA sequencing and *in situ* hybridization chain reaction (HCR). We show that conserved developmental regulators, including Hox and Wnt pathway genes, exhibit region-specific expression corresponding to distinct budding zones and zooid distributions. These results indicate that canonical axial patterning systems are deployed at the level of the colony axis. Our findings demonstrate that developmental gene networks classically associated with anterior-posterior patterning can operate at a higher level of biological organization, providing a mechanistic framework for the evolution of integrated, superorganism-like body plans in colonial animals.

## Background

Colonial animals can achieve functional integration by spatially organizing specialized modules along a shared axis, in some cases functioning as superorganism-like entities^1,2^. Siphonophores (Cnidaria; Hydrozoa) exemplify this phenomenon, forming highly integrated colonies in which genetically identical zooids are precisely arranged along an elongating stem to perform distinct functions, including locomotion, feeding, and reproduction^1,3-5^ (**Figure 1**).

**Figure 1.**
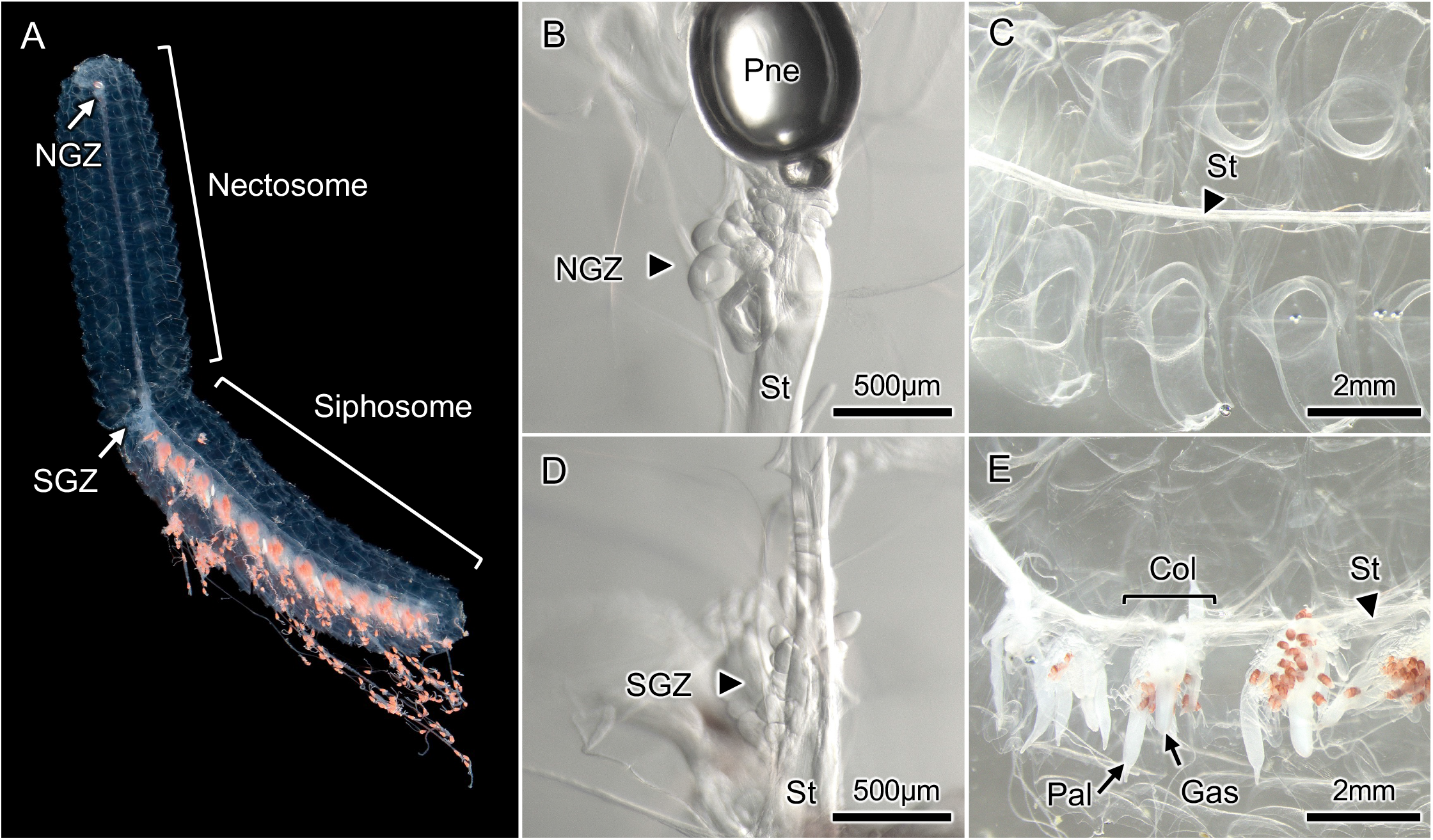
Overall colony organization of the siphonophore *Agalma okenii*. (A) Overview image of the colony. The colony is divided into the nectosome and siphosome, each containing a distinct budding region: the nectosomal growth zone (NGZ) and the siphosomal growth zone (SGZ), respectively. (B) Enlarged image of the nectosomal growth zone (NGZ). A pneumatophore (Pne), which contributes to buoyancy regulation of the colony, is located at the apical region of the stem (St), and small, early-stage swimming zooids (nectophores) are visible immediately below it. (C) Enlarged image of the siphosomal growth zone (SGZ). Multiple zooid types bud from the stem (St), including feeding gastrozooids (Gas), digestive palpons (Pal), and bracts that contribute to colony structure. (D) Enlarged image of the mid-nectosome. Well-developed nectophores are arranged around the central stem (St). (E) Enlarged image of the mid-siphosome. Gastrozooids (Gas), palpons (Pal), and gonozooids are arranged along the stem, with bracts surrounding these zooids.

All zooids arise from two spatially separated budding zones along the stem: the nectosomal and siphosomal growth zones^4,6-8^. In the nectosomal growth zone (NGZ), only nectophores (swimming zooids) are produced, whereas the siphosomal growth zone (SGZ) generates multiple functionally specialized zooid types, including gastrozooids (feeding zooids), palpons (digestive and storage zooids), gonodendra (reproductive units), and bracts, which provide structural support and protection. This strict association between growth zones and zooid types suggests that positional information along the stem, particularly within each growth zone, governs both zooid identity and spatial organization (**Figure 1**).

Although previous transcriptomic studies have identified genes associated with the morphology and function of individual zooids^9^, how axial information is encoded along the stem itself has remained unclear. To address this question, we examined gene expression patterns along the stem of the siphonophore *Agalma okenii*, with particular focus on the two budding zones where positional cues are expected to be established and maintained. However, how such axial information is encoded at the colony level remains unknown.

## Results and discussion

### Transcriptomic analysis of distinct stem regions

To characterize gene expression profiles associated with each budding zone, we dissected stem regions containing either the nectosomal growth zone or siphosomal growth zone and further separated the budding zone from adjacent non-budding stem tissue. After removing zooids and associated structures such as bracts, we extracted total RNA from each dissected region and performed RNA sequencing–based transcriptomic analyses after *de novo* transcriptome assembly (**Figure S1, Table S1**).

Principal component analysis separated budding zones (nectosomal growth zone and siphosomal growth zone) from non-budding stem regions along PC1, while PC2 captured variation along the anterior–posterior axis (**Figure S2**). Differential expression analyses and GO (gene ontology) term enrichment analyses revealed strong regionalization of gene expression (**Figure S3–4, Table S2–4**). Genes associated with cell proliferation were enriched in both growth zones (**Figure S4C, E**), consistent with their role as sites of active zooid production. Stem cell markers (e.g., *Piwi*) were highly expressed in the both zones, whereas germ cell markers (e.g., *Dazl*) were enriched specifically in the siphosomal growth zone (**Table S2–4**), reflecting its role in reproductive zooid formation (**Figure 1**). Moreover, gene expression associated with collagen and muscle development is elevated in the nectosomal growth zone, suggesting its involvement in the formation of locomotory organs that enable movement (**Table S3**).

Previous studies suggested that conserved developmental regulators, including Hox genes and Wnt pathway components, contribute to colony formation and zooid differentiation^9-11^. Based on this view, we focused in particular on Hox genes involved in oral–aboral axis patterning and components of the canonical Wnt signaling pathway, key regulators of axial specification in cnidarians. Many of these genes exhibited budding zone–specific expression (**Figure 2A**), suggesting that conserved axial patterning modules may be redeployed in the specification of growth zone. Genes such as *AntHox6* and *Wnt3*, showed higher expression in the siphosomal growth zone, whereas genes including *AntHox1* and *Dkk1* (*Dickkopf-1*), were more highly expressed in the nectosomal growth zone (**Figure 2A**). In cnidarians, these genes are associated with the oral–aboral patterning during development. In particular, *AntHox6* and *Wnt3* are involved in oral specification, whereas *AntHox1* and *Dkk1* contribute to aboral specification and patterning^12^. The expression dynamics observed here reveal spatially segregated oral and aboral gene expression across distinct budding zones along the stem, suggesting that the siphonophore colony exhibits axial patterning reminiscent of that of a single individual.

**Figure 2.**
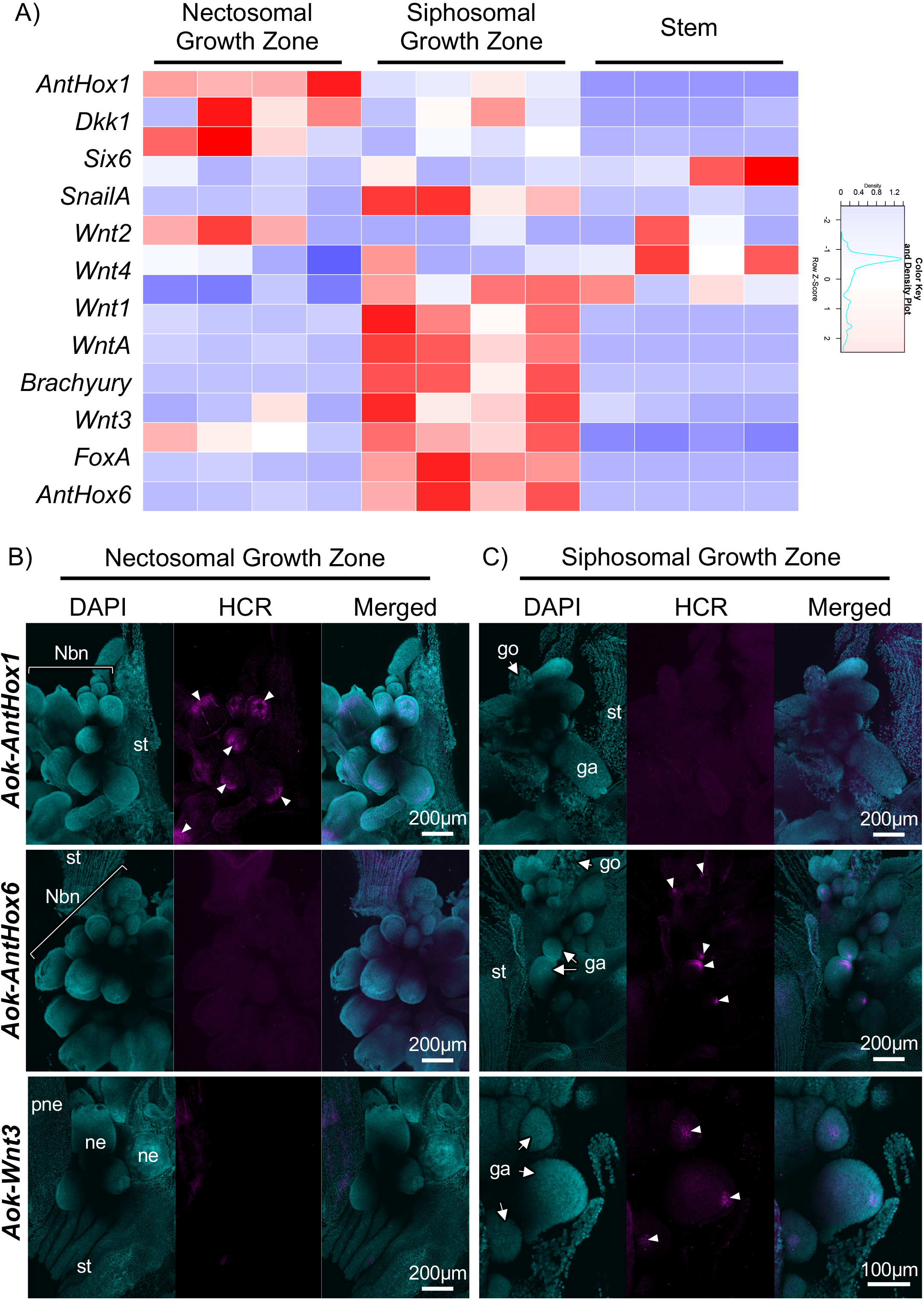
Growth zone specific expressions of axial patterning gene. (A) Heatmap showing the expression dynamics of genes involved in metazoan axial patterning and developmental regulation, including Hox and Wnt pathway–related genes, across the different stem regions. Expression values are shown as TPM-based Z-scores. (B–C) Localization of representative Hox and Wnt pathway–related gene expression revealed by *in situ* hybridization chain reaction (HCR). Expression patterns of *AntHox1, AntHox6*, and *Wnt3* are shown in (B) the nectosomal growth zone (NGZ) and (C) the siphosomal growth zone (SGZ). *AntHox1* is expressed within the newly budding nectophore (nbn), whereas *AntHox6* and *Wnt3* are expressed on the oral side of gastrozooids (ga). go indicates the gonodendron. Magenta fluorescence indicates the localization of target gene transcripts, and cyan indicates nuclei stained with DAPI.

**Figure 3.**
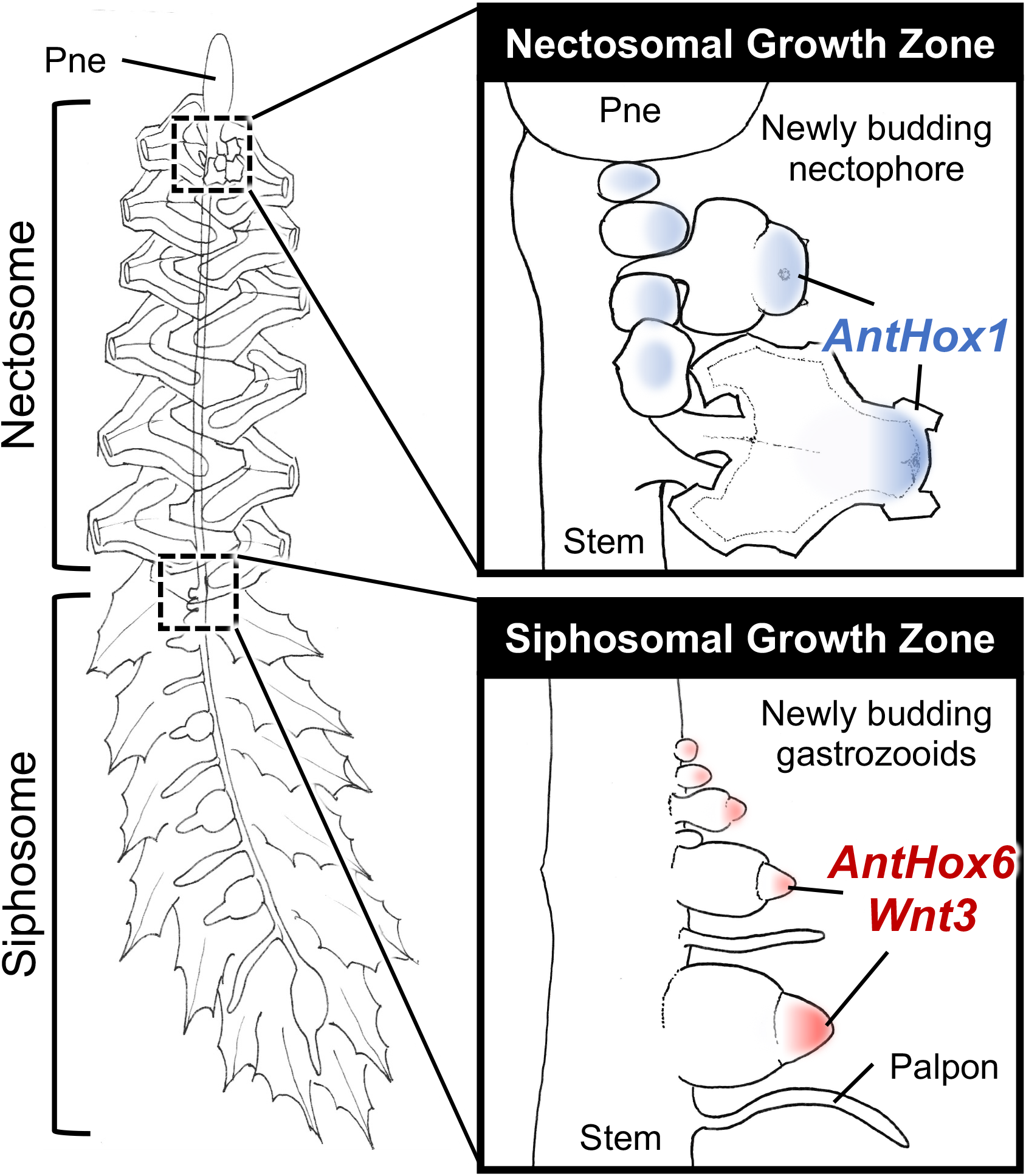
Conserved axial patterning gene networks underlying colony-level organization in siphonophores. Hox genes and components of the Wnt signaling pathway, which regulate anterior–posterior axis specification in bilaterians and oral–aboral patterning in cnidarians, are differentially deployed along the siphonophore stem. These conserved developmental regulators operate at the colony level to generate spatially ordered differentiation and precise placement of functionally specialized zooids. In this model, colony organization emerges from the extension of canonical axial patterning systems across the entire stem, rather than from independent patterning processes within individual zooids.

### Spatial deployment of axial patterning genes

To resolve the spatial organization of regionally expressed genes, we focused on *AntHox1, AntHox6*, and *Wnt3*, which showed robust and consistent SGZ- or NGZ-specific expression across transcriptomic datasets (**Figure 2A**). Homologs of these genes function as master regulators of oral–aboral axis specification in *Nematostella vectensis*^12^, implicating them in axial patterning. We therefore examined their spatial expression using Hybridization Chain Reaction (HCR).

Consistent with transcriptomic results, *AntHox1*, enriched in the nectosomal growth zone, was detected in endodermal tissues of early budding nectosomes, i.e., swimming zooids (**Figure 2B**) and was absent from the siphosomal growth zone (**Figure 2C**). In contrast, *AntHox6* and *Wnt3*, both enriched in the siphosomal growth zone, showed HCR signals restricted to this region (**Figure 2C**). These signals were particularly prominent at the oral tip of developing gastrozooids, indicating spatially confined expression associated with zooid differentiation.

### Colony-level axial patterning as a developmental system

Hox genes and the Wnt signaling pathway are central regulators of anterior–posterior axis specification in animals and also contribute to oral–aboral patterning in cnidarians^12-15^. Our results demonstrate that these conserved axial patterning gene networks operate at the level of the siphonophore cology, generating spatially ordered differentiation of zooids along the stem.

In this framework, zooid placement arises from the extension of canonical developmental patterning systems from individual-level axis formation to the scale of the entire colony, such that positional information is established and interpreted across the colony axis rather than independently within individual zooids. Thus, the formation of a functionally integrated, organism-like colony relies on developmental mechanisms fundamentally similar to those governing axis formation in individual development.

Classical descriptions of siphonophore development indicate that the colony originates from a primary gastrozooid, from which the stem elongates aborally and produces additional zooids through budding^8,16-18^. This developmental trajectory implies that the colony axis derives from the oral–aboral axis of the founding individual, i.e., primary gastrozooid (**Figure S5**). The mature siphonophore colony may therefore represent a higher-order structure organized around an original individual (i.e., primary gastrozooid) axis, with zooids budding along the stem and some becoming organ-like units.

Haeckel (1866) treated siphonophore colonies as higher-order individuals composed of polymorphic but physiologically integrated parts^19^. However, as shown here, the presence of a unified axial patterning system along the stem suggests that the siphonophore colony may instead be understood as a single morphogenetic individual. In this framework, zooids can be interpreted as modular units integrated into a higher-level developmental system. This interpretation is consistent with, but extends beyond, earlier perspectives such as that of Garstang (1938)^20^, by showing that colony-level individuality is underpinned by conserved developmental gene networks classically associated with axis formation in single organisms. Our results thus integrate these interpretations, showing that siphonophore colonies function as single morphogenetic individuals (cormi) in which zooids act as modular, organ-like units.

Future studies integrating temporal analyses of gene expression with functional analysis during early colony development will be essential to test this hypothesis and clarify how individual-level developmental programs are extended to generate colony-level body plans. More broadly, our findings demonstrate that canonical developmental gene networks can be co-opted to operate across hierarchical levels of biological organization, providing a mechanistic basis for the evolution of superorganism-like body plans.

## Materials and Methods

### Sample collection and morphological observation

Samples of *A. okenii* were collected from rocky shores, beaches, and offshore waters around Misaki Marine Biological Station in Kanagawa Prefecture, Japan, between December and February from 2020 to 2024. Specimens were collected either by snorkeling, using zipper bags to capture colonies underwater, or from a boat by scooping swimming colonies from the sea surface with a plastic dipper. Specimens were anesthetized in 3.5% MgCl□, immediately dissected using fine tweezers under a stereomicroscope (SZX16; Olympus, Tokyo, Japan), and either fixed in 4% paraformaldehyde (PFA) or processed directly for RNA extraction.

### RNA sequencing and differentially expressed gene analysis

To characterize gene expression profiles across netsomal growth zone, siphosomal growth zone and other region in stem, RNA sequencing was performed. Total RNAs were extracted from the dissected region using RNAiso Plus (Takara Bio, Shiga, Japan). After extraction, samples were treated with DNaseI (Thermo Fisher Scientific, Waltham, MA, USA) and Agencourt AMPure XP beads (Beckman Coulter, Brea, CA, USA) for purification. Complementary DNA libraries were constructed using TruSeq RNA Sample Preparation Kits version 2 (Illumina Inc., San Diego, CA) and sequenced by HiSeq X (Illumina Inc.).

After RNA sequencing, quality of all obtained data was checked by the FastQC^21^ version 0.11.9, and adaptors and low quality sequences were trimmed using Trimmomatic^22^ version 0.39. All of the filtered sequencing reads were pooled and subjected to de novo assembly using Trinity^23^ version 2.13.1 with default parameter. Protein-coding sequences were estimated using TransDecoder version 5.5.0. To evaluate assembly completeness, Assembled contigs and estimated peptide sequences were subjected to BUSCO^24^ version 6.0.0 using the single copy ortholog set of metazoa_odb12 and cnidaria_odb12. Orthologs in the focal species were annotated based on BLASTP search against *Hydra vulgaris* RefSeq peptide sequences, using an e-value threshold of < 1.0e-4. To remove redundant sequences (isoforms) in RefSeq peptides, hydra’s longest isoform from each gene was obtained from the NCBI RefSeq genome (GCF_038396675.1) using the “retrieve_longest_isoforms” function in the R package orthologr^25^ version 0.4.2. Also, to characterize focal developmental regulators (listed in **Figure 2A**) carefully, reciprocal blast best hit analysis using BLASTP search with Sequenceserver^26^ version 2.0.0 were preformed against peptide sequences of *Nematostella vectiensis* ^12^, using e-value threshold of < 1.0e-500.

For gene expression analyses, filtered reads were then mapped to Trinity assembled contigs using Salmon^27^ version 1.0.0 and obtained TPM values. To summarize gene expression patterns across samples, genes with TPM ≥ 1 in all 12 samples were extracted, expression values were normalized as log_10_(TPM + 1), and then principal component analysis was performed using the top 2,000 most variable genes. Differentially expressed genes (DEGs) were identified using a generalized linear model (GLM) framework with quasi-likelihood F tests in an R package edgeR^28^ version 4.8.2. The false discovery rate (FDR) was controlled at 5%. Genes with adjusted *P*-value < 0.05 and |log2 Fold Change| > 1 were considered DEGs.

Gene ontology (GO) term enrichment analyses were performed using clusterProfiler^29^ version 4.18.4. Using NCBI gene2go and gene2refseq (last accessed: 2026/02/24), GO terms assigned to *H. vulgaris* genes were transferred to the corresponding sequences of the focal species based on BLASTP hits (see above). Then, custom gene ontology databases were built using an R package, AnnotationForge version 1.52.0 (https://doi.org/10.18129/B9.bioc.AnnotationForge). GO term enrichment analyses were performed using “enrichGO” and “simplify” functions in clusterProfiler with adjusted *P*-values < 0.05 as thresholds. All statistical analyses were performed using R 4.2.2 (https://www.r-project.org).

### In situ Hybridization chain reaction (HCR)

To investigate the precise localization of developmental regulation genes, gene expression in zooid growth zones was examined using whole-mount HCR. Samples were fixed overnight in 4% PFA in PBS and stored in methanol at -80℃. Before experiments, samples were dissected into two or three segments to facilitate probe penetration and washed with 1x PBST in room temperature. Experiments were performed using the ISH Palette™ series kits (Nepagene, Chiba, Japan), which includes the necessary buffers and harpin DNA, largely following a previously described protocol^30^. Nuclei were counterstained with 2 µg/mL of 4’,6-diamidino-2-phenylindole (DAPI; Sigma, St. Louis, MO, USA) for one hour at room temperature, after completion of the amplification reaction with hairpin DNA (IPL-G-S23) labeled with Alexa fluor 488 and subsequent wash with PBST were completed. To specifically label the genes of interest, paired oligonucleotide DNA probes containing an initiator sequence for amplification of hairpin DNA were custom designed by Nepagene and synthesized by Fasmac (Kanagawa, Japan) (**Table S5**). All samples were observed using a confocal microscope (FV3000; Olympus).

## Acknowledgements

We would like to express our gratitude to Kazuhisa Hori, Mitsugu Kitada, Aya Adachi, Gaku Yamamoto, Michiyo Kawabata, Izumi Komori, Megumi Kohtsuka, Yohei Otomo, Ryosuke Kimbara, Soma Chiyoda, Naoki Kanai, Yukio Kurihara, Hiromi Kurihara and the Miura Fishery Cooperative Association for their help with specimen sampling. We would also like to offer special thanks to Soma Chiyoda for providing photographs (**Fig. 1A**). This work was supported by a Grant-in-Aid for Research Activity Start-up (No. 22K20662) from Japan Society for the Promotion of Science, Research Institute of Marine Invertebrates and Sasakawa Grants for Science Fellows (SGSF) for K.O..

## Author Contributions

K.O., S.K. and T.M. conceptualized and designed the study. K.O. and H.K. performed field sampling. K.O. and A.Y. performed transcriptome analysis. K.O. performed morphological observations and gene expression analysis with HCR. All authors wrote and approved the final version of the manuscript.

## Additional Information Competing interests

The authors declare no competing interests.

## Data availability

Raw sequence reads and assembled contigs generated using Trinity have been deposited in NCBI/DDBJ/EMBL database under the Bioproject accession number PRJDB40757.

## Ethics Declarations

No ethical approval is required.

---

**Table S1**. Statistics of *de novo* assembled contigs of *A. okenii*.

**Table S2**. Differentially expressed genes between the siphosomal growth zone (SGZ) and the nectosomal growth zone (NGZ). “Up”-regulated genes were highly expressed in SGZ, whereas “Down”-regulated genes were highly expressed in NGZ. Differentially expressed genes (DEGs) were defined as those with adjusted *P*-value (p.adj) < 0.05 and |log2FoldChange (log2FC)| > 1.

**Table S3**. Differentially expressed genes between the nectosomal growth zone (NGZ) and stem part. “Up”-regulated genes were highly expressed in NGZ, whereas “Down”-regulated genes were highly expressed in stem part. Differentially expressed genes (DEGs) were defined as those with adjusted *P*-value (p.adj) < 0.05 and |log2FoldChange (log2FC)| > 1.

**Table S4**. Differentially expressed genes between the siphosomal growth zone (SGZ) and stem part. “Up”-regulated genes were highly expressed in SGZ, whereas “Down”-regulated genes were highly expressed in stem part. Differentially expressed genes (DEGs) were defined as those with adjusted *P*-value (p.adj) < 0.05 and |log2FoldChange (log2FC)| > 1.

**Table S5**. List of probe sets used in HCR.

**Figure S1**. Completeness of *de novo* assembled contigs of *A. okenii*. (A) BUSCO scores of the assembled contigs against odb12_metazoa and odb12_cnidaria (B) BUSCO scores of the estimated peptide sequences against odb12_metazoa and odb12_cnidaria.

**Figure S2**. Principal component analysis of transcriptome data. PCA were performed using top 2,000 most variable genes (TPM values) after log_10_ normalization. red: the siphosomal growth zone (SGZ); blue: the nectosomal growth zone (NGZ); and green: the stem part.

**Figure S3**. Results of differential gene expression analyses. (A–C) MA plot for all three pairwise comparisons. Differentially expressed genes (DEGs) were defined as those with adjusted *P*-value < 0.05 and |log2FoldChange| > 1, and are highlighted as follows: red, highly expressed at the siphosomal growth zone (SGZ); blue, highly expressed at the nectosomal growth zone (NGZ); and green, highly expressed at the stem. (D) Venn diagram of the numbers of shared DEGs across three comparisons.

**Figure S4**. Top 10 enriched gene ontology terms of biological functions in DEGs of each comparisons. For each category, the ten GO terms with the lowest adjusted P values are shown. Enriched GO terms are shown for DEGs with higher expression in (A) SGZ compared with NGZ, (B) NGZ compared with SGZ, (C) NGZ compared with stem, (D) stem compared with NGZ, (E) SGZ compared with stem, (F) stem compared with NGZ.

**Figure S5**. Colony development and axis formation in siphonophores. (A) The colony axis is established along the oral–aboral axis of the primary gastrozooid, from whose aboral side the stem elongates and produces additional zooids by budding. Arrows indicate the body axis, with the red arrowhead marking the oral side. Blue bars protruding from the stem represent the axial information of nectophores, specified by aboral patterning factors. (B) A hypothetical evolutionary scenario for the emergence of highly sophisticated colonies such as those of siphonophores. In ancestral colonial hydrozoans, oral–aboral axis patterning is established independently within individual zooids. In siphonophores, this patterning system is integrated during development into a colony-level axis, as shown in (A), resulting in a highly coordinated, organism-like body plan.

