## Supplementary figures and images for "Axial Patterning Beyond the Individual: Colony-level Organization in a Siphonophore Colony"

### Supplemental Figure 1

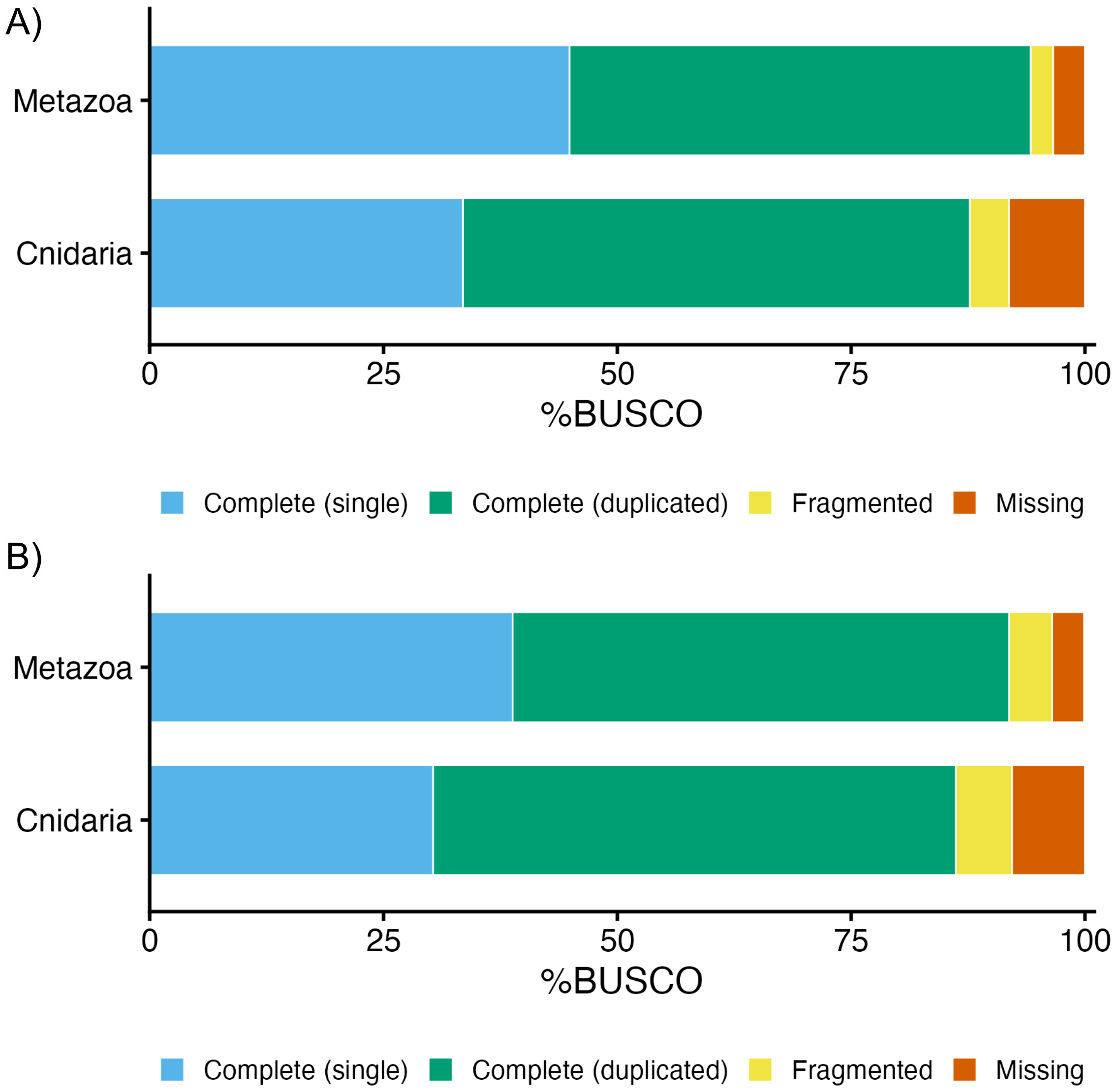

### Supplemental Figure 2

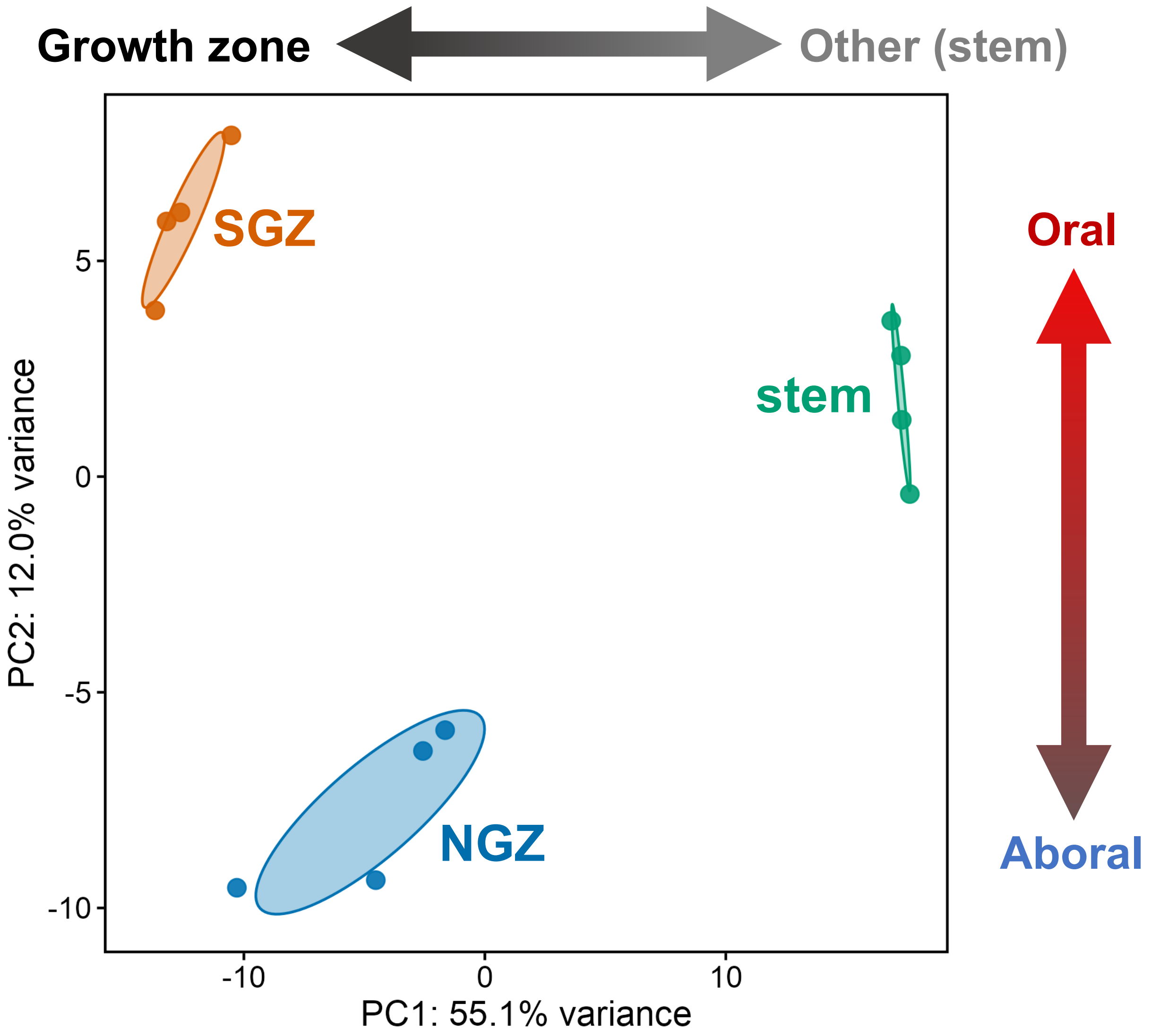

### Supplemental Figure 3

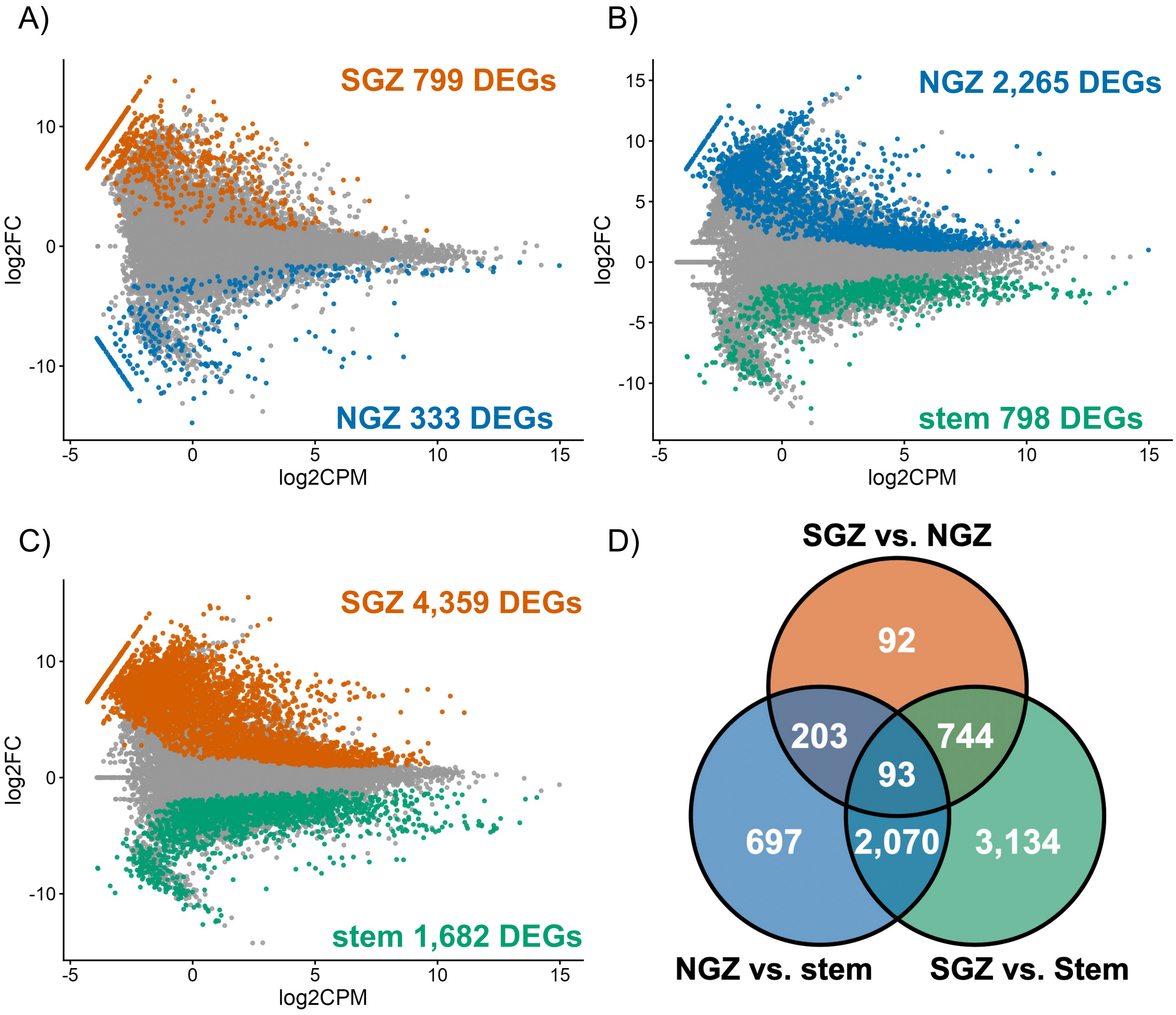

### Supplemental Figure 4

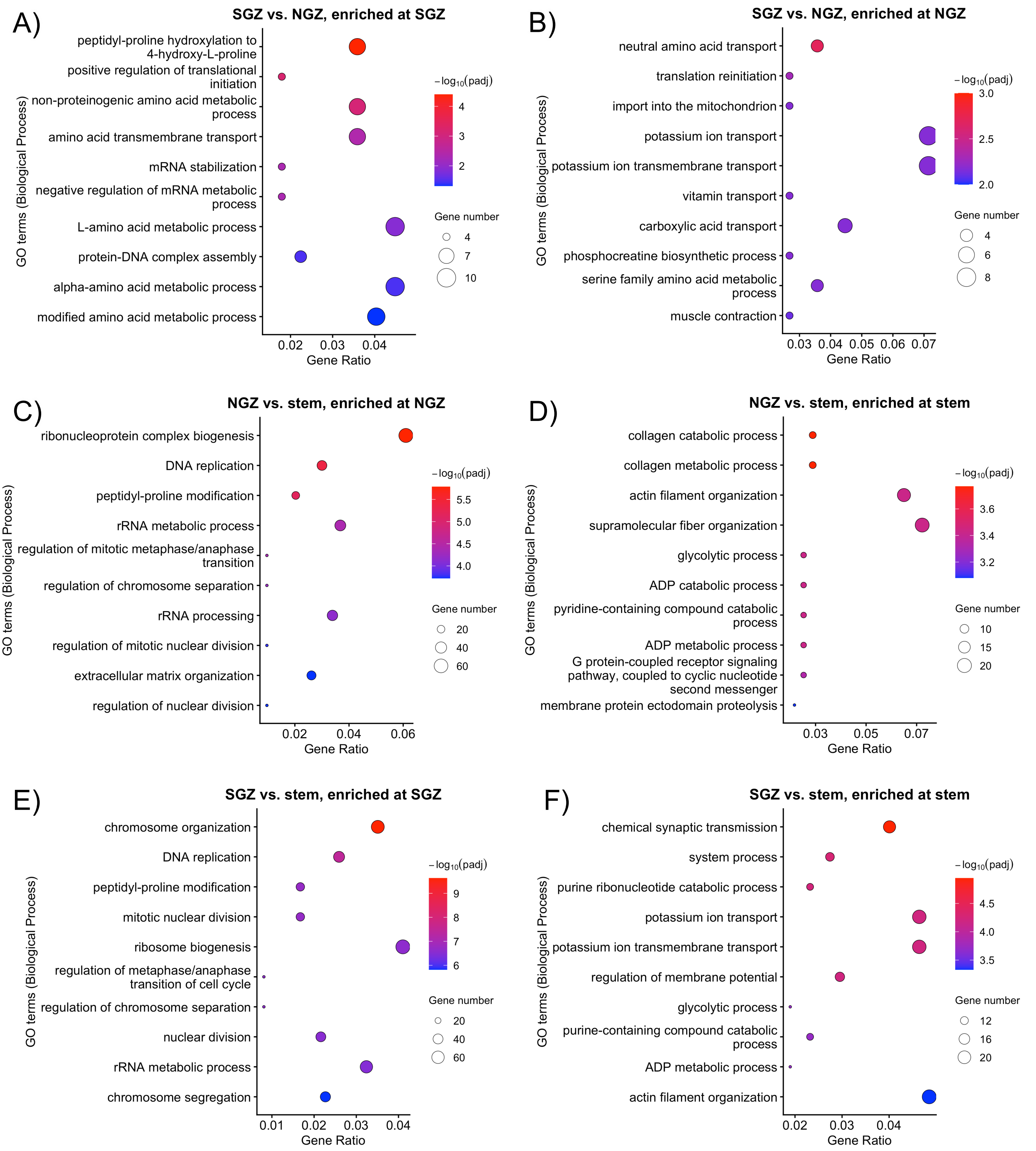

### Supplemental Figure 5

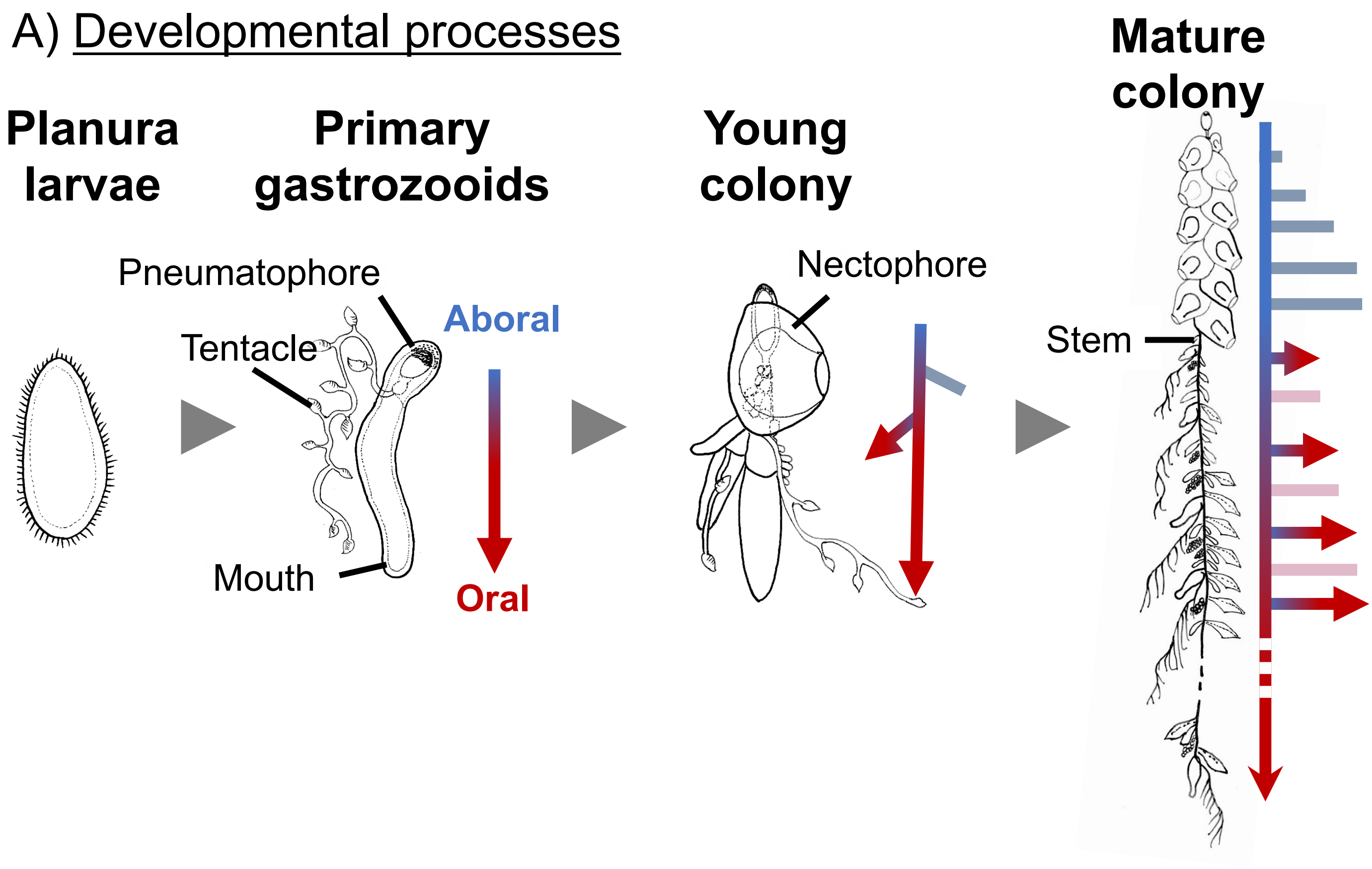

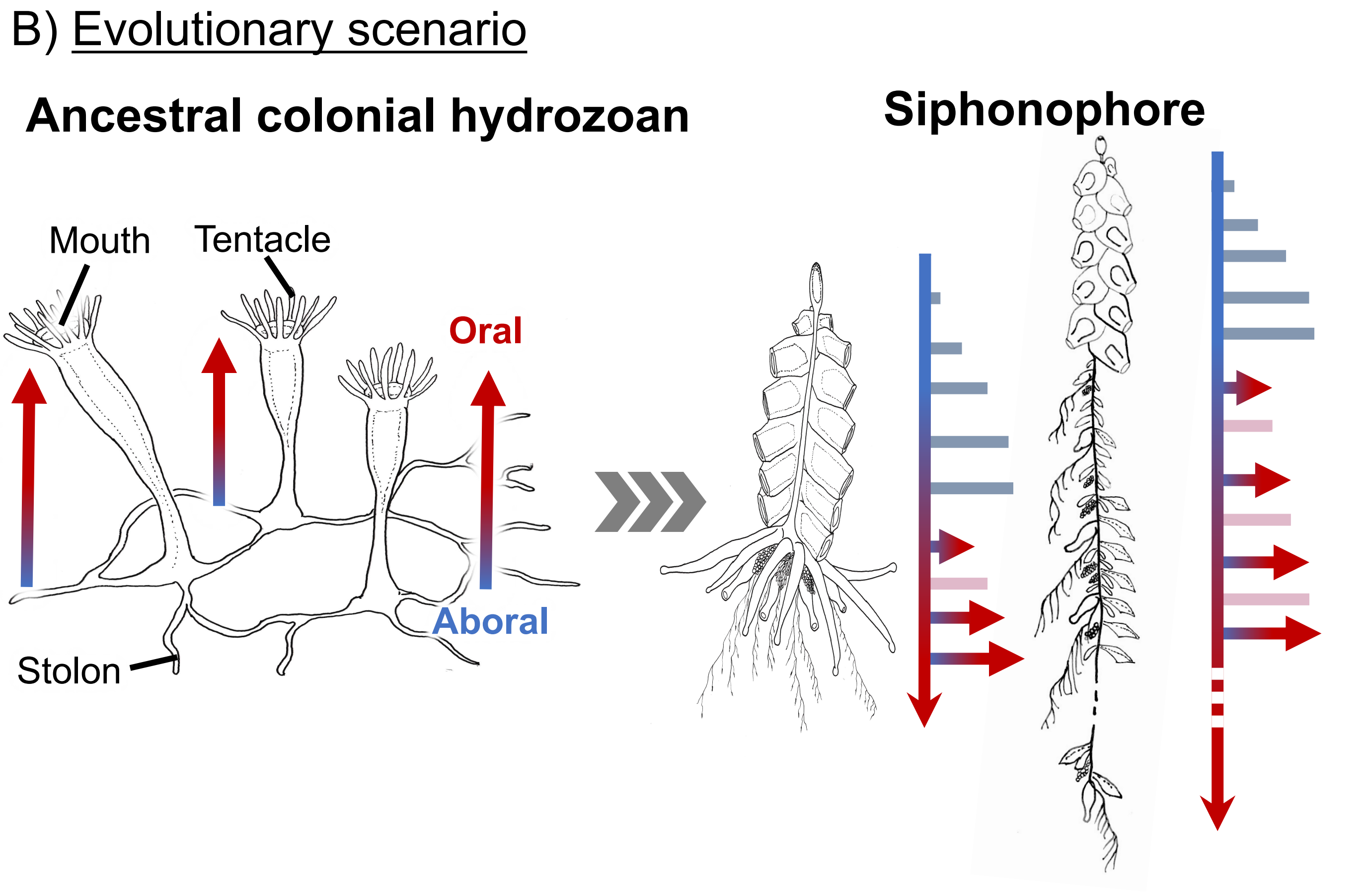
